# Exact distribution of threshold-crossing times for protein concentrations: Implication for biological timekeeping

**DOI:** 10.1101/2021.08.20.457050

**Authors:** Krishna Rijal, Ashok Prasad, Abhyudai Singh, Dibyendu Das

## Abstract

Stochastic transcription and translation dynamics of protein accumulation up to some concentration threshold sets the timing of many cellular physiological processes. Here we obtain the exact distribution of first threshold-crossing times of protein concentration, in either Laplace or time domain, and its associated cumulants: mean, variance and skewness. The distribution is asymmetric and its skewness non-monotonically varies with the threshold. We study lysis times of E-coli cells for holin gene mutants of bacteriophage-λ and find a good match with theory. Mutants requiring higher holin thresholds show more skewed lysis time distributions as predicted.

---

Molecular biological processes associated with changes in cell state are controlled by changes in gene expression, a complex stochastic process involving transcription of a gene into an RNA molecule and its subsequent translation into a protein. Intrinsic noise in transcription and translation leads to stochastically varying abundance of mRNA and proteins within cells, even when genetically identical [1–3]. Experimental studies have directly probed fluctuations of protein levels across cells [4, 5]. Theoretical models assuming specific promoter configurations have solved for the co-efficient of variance *CV*^2^ of the mRNA and protein numbers, as well as their full steady state distributions [6–10]. Stochastic gene expression has been shown to be a fundamental property of living cells, affecting critical physiological processes of biological and biomedical importance [6].

The dynamics of downstream processes governed by the synthesis of a protein requires its accumulation to some minimum concentration threshold, eg. transcription factors with sigmoidal Hill kinetics [11]. In such cases the timing of the downstream process is governed by the time at which the threshold concentration is reached for the first time, i.e. the *First Passage Time* (FPT) [12, 13] of the stochastic gene expression process. The most well studied example of timing control by stochastic protein accumulation is probably lysis of lambda phage-infected *E. Coli*. Here the protein Holin self-assembles on the bacterial membrane and punctures it after its concentration crosses a threshold, causing the cell to ultimately lyse or burst and release the newly formed viral particles [14, 15]. While previous work has studied the timing of lysis and its relation with viral fitness, [15–17], later studies have highlighted the distribution of lysis timing and demonstrated connections with the first passage time [18–21]. Fluctuations in lysis times lead to variations in viral burst sizes affecting both viral population fitness [22] as well as health of the host. Genetic mutations of Holin have been shown to regulate the stochasticity in lysis times in the *λ*-variants [18, 23, 24].

First passage times based mechanisms may be quite common in cells since recent work suggests that it operates in the expression of *mig1* in C. Elegans [25], and *FtsZ* expression in bacterial cell divisions [26]. First passage times are also relevant for other phenomena such as RNA polymerase backtracking and cleavage [27], first binding of proteins to sites on DNA [28], capture of kinetochores [29] and, more abstractly, estimating characteristics of energy landscapes [30]. Statistics of first passage times (FPT) have been of great theoretical interest and obtaining analytical expressions for their distribution (FPTD) is generally quite challenging [31]. In previous work we derived analytical expressions for the FPTD of the absolute number of any molecule generated through geometrically distributed burst kinetics to reach a threshold[32]. However for biological applications the relevant variable is not the absolute number but the concentration, since cellular volume is not constant either across cells or even in a single cell over time. Previously only approximate formulae existed for the moments of the threshold-crossing FPTD [20, 24], and the distribution itself was unknown. Here we derive exact analytical expressions (as well as systematic approximations), for the FPTD of molecular concentrations and its moments, and apply them to experimental data on the distribution of lysis times.

A theoretical framework for the stochastic kinetics of protein synthesis has already been developed under the assumptions of short-lived mRNA and long-lived proteins. The exact steady-state distributions of the discrete protein number (*n*) has been derived previously [8] and was shown to follow a negative binomial distribution, while the continuous protein concentration *c* = *n/V* (with *V* being cellular volume) was shown to follow the Gamma distribution [7]. While the *forward* continuous Master equation was suitable to study the protein concentration *c* [7], the corresponding *backward* Master equation [33] is more convenient for calculating the statistics of the FPT to reach the threshold concentration *X*. Given an *initial c* = *x* < *X*, the survival probability *S*(*X, x, t*) that *c* survives reaching the threshold *X* through time *t*, satisfies the backward continuous Master equation:

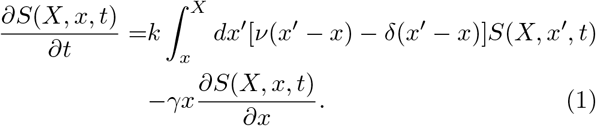

Here the the initial condition is *S*(*X, x*, 0) = 1 and boundary condition *S*(*X, x* = *X, t*) = 0, and the rate of protein production *kν* is assumed to be proportional to the experimentally known protein burst size distribution 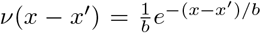 with mean burst concentration *b* [5, 34]. The rate *γx* of concentration decay expresses the joint effect of protein degradation and cell growth. The FPTD for the *first* threshold crossing (*x* ≥ *X*) in time *t* is obtained from *S*(*X, x, t*) as: *f* (*X, x, t*) = −*∂S*(*X, x, t*)*/∂t*. To solve Eq. (1) we convert the integro-differential equation into a partial differential equation and take the Laplace Transform, 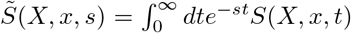 leading to a differential equation for 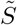 as a function of the (scaled) initial concentration 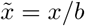 (details in S2 of SI [35]):

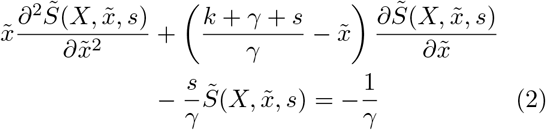

The homogeneous part of the above equation is a confluent Hypergeometric equation [36]. Using the boundary condition and the fact that 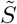 is finite as *x* → 0, the solution in terms of the confluent Hypergeometric function 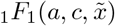 [36] is as follows (see S3 of SI [35]):

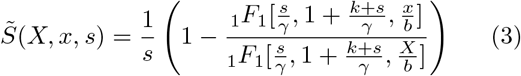

Since 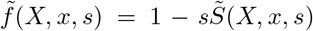, the desired exact FPTD in Laplace space for any *γ* and any initial protein concentration *x* > 0 is:

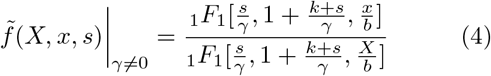

The above calculation (for *x* > 0) is applicable to the special case *x* → 0 of our interest. The case of exactly *x* = 0 requires a separate treatment, but is numerically identical to *x* → 0 as expected (details in S3 of SI [35]).

For vanishing decay constant (*γ* → 0), the Eq.(4) simplifies to 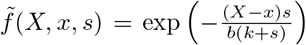 which is analytically invertible and gives the exact FPTD in the time domain (see S4 of SI [35]):

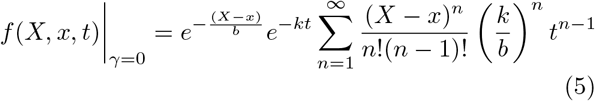

Note that for *γ* = 0 the result is a function of the difference between the threshold and initial concentrations (*X* − *x*), as expected from the translational symmetry in Eq. (1).

In general for *γ* ≠ 0, the Eq.(4) may be inverted numerically using Mathematica [37]. Finally we can also derive the exact expression of the first few moments, which are useful when comparing with empirical distributions. From Eq.(4), the *n*-th moment 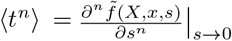,. Defining 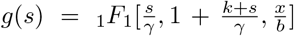, 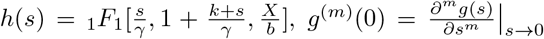 and 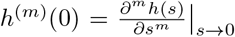, the first three moments are derived analytically exactly (see S5 of SI [35]):

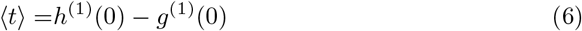

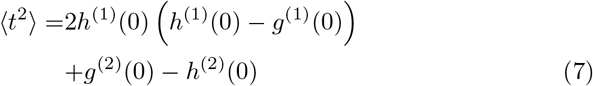

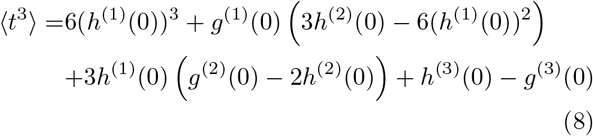

Note that 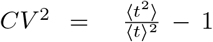 and the Skewness = 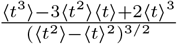 follow from the above expressions.

Just like the distribution in Eq.(4), the quantities *g*^(1)^(0), *g*^(2)^(0), *g*^(3)^(0), *h*^(1)^(0), *h*^(2)^(0) and *h*^(3)^(0) (see S5 of SI [35]) depend on the four parameters (*x/b*), *k, γ* and (*X/b*).

## Statistics of Lysis Times

In order to match our results with experimental data, we used the raw data of Ref. [24]. Briefly, site-directed mutagenesis was used to generate a library of mutations in the S105 holin allele, each of which differed from the parent allele by one or two amino acid substitutions. These mutated sequences were then used to generate a library of lysogenic lambda phages, each carrying a slightly different holin gene. These viruses were used to infect E. coli cells and lysis was thermally induced and measured at the single cell level for between 91-174 cells per strain.

We estimate some required parameters as follows. Holins degrade slowly hence the decay of *x* is mostly due to cell growth, with doubling time of roughly 40 mins. Hence we choose *γ* = ln(2)/40 min^−1^. We choose *x/b* = 0.01 to represent *x* → 0, the vanishingly small initial protein concentration. Next we numerically eliminate the parameter (*X/b*) between the expressions of *CV*^2^ and mean FPT ⟨*t*⟩ (see discussion S6 in SI [35]), such that the theoretical curve of *CV*^2^ versus ⟨*t*⟩ gets fixed by just one fitting parameter i.e. *k*. The best fit of the theory to the experimental data for the 20 mutants is shown in Fig.1, and yields the fitted value *k* = 4.5 *min*^−1^. The *CV*^2^ curve has a minimum at mean FPT around *t*_*m*_ ∼ 40 mins.

**FIG. 1:**
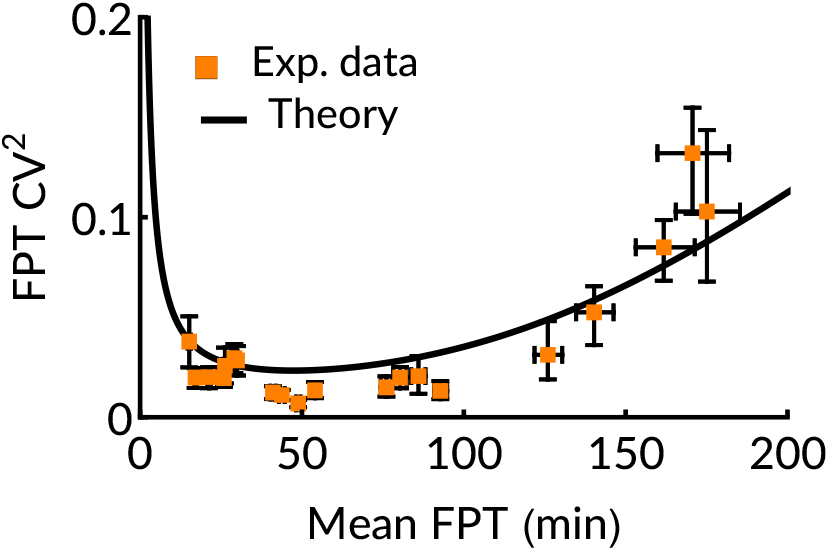
*CV*^2^ against ⟨*t*⟩ for 20 mutants (symbols) and exact theory (solid line) with best fit parameter *k* = 4.5 *min*^−1^. Error bars are 90% confidence intervals, obtained after bootstrapping (1000 replicates). Here *γ* = ln(2)/40 min^−1^ and 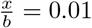.

While we expect the mutants to have roughly same (*x/b*), *k* and *γ* values (as given above), their mean FTP differs as does the threshold (*X/b*). We fix (*X/b*) for every mutant by matching the theoretical mean Eq. (6) with the experimental average from data, for that mutant. Then we obtain the full theoretical FPTD (by inverting Eq.(4) [37]) and plot against the experimental distribution to check how well they match. This is shown in Fig.2 for two cases — mutant-1 (JD405) and mutant-2 (JD426) (see S1 of SI [35] and [24]) have mean FPT smaller and larger than *t*_*m*_ respectively.

**FIG. 2:**
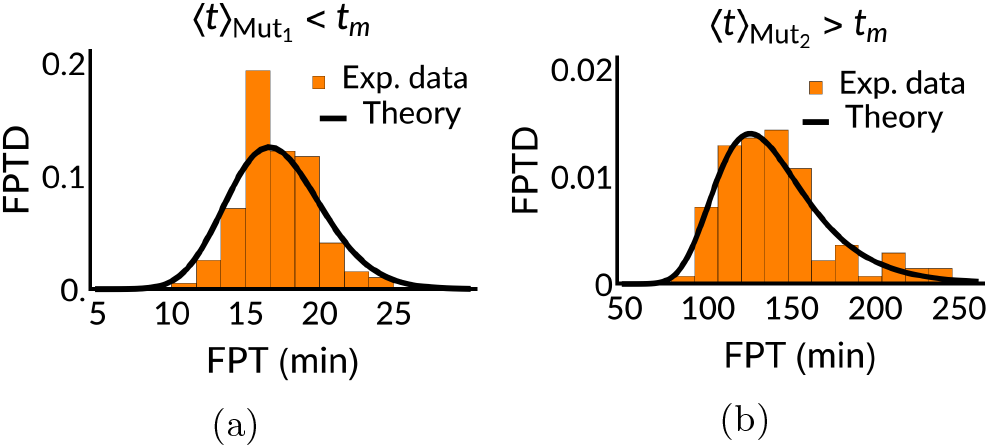
The FPTD from experiment and theory for (a) mutant-1 with ⟨*t*⟩ = 17.1 mins and (b) mutant-2 with ⟨*t*⟩ = 140.3 mins. Note *t*_*m*_ ∼ 40 mins.

The data and theoretical curves (in Fig.2) both suggest that FPTD is non-Gaussian and is skewed to the right. We explicitly study the variation of the Skewness of FPTD for the mutants in Fig.3 — it shows a non-monotonic behaviour just like the *CV*^2^ with a minimum around ∼35 mins close to *t*_*m*_ mentioned above. Thus mutants with increasingly larger mean FPT, have increasingly asymmetric FPTD. Note that the distributions have asymptotic (large *t*) exponential tails ∼ exp(−*t/τ*_*c*_), with characteristic times *τ*_*c*_ being related to the smallest pole *s*_*\**_ = −1*/τ*_*c*_ of 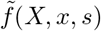 in Eq.(4). A plot of *τ*_*c*_/ ⟨*t*⟩ against mean FPT shows a similar non-monotonic curve as *CV*^2^ and Skewness (see S7 in SI [35]).

**FIG. 3:**
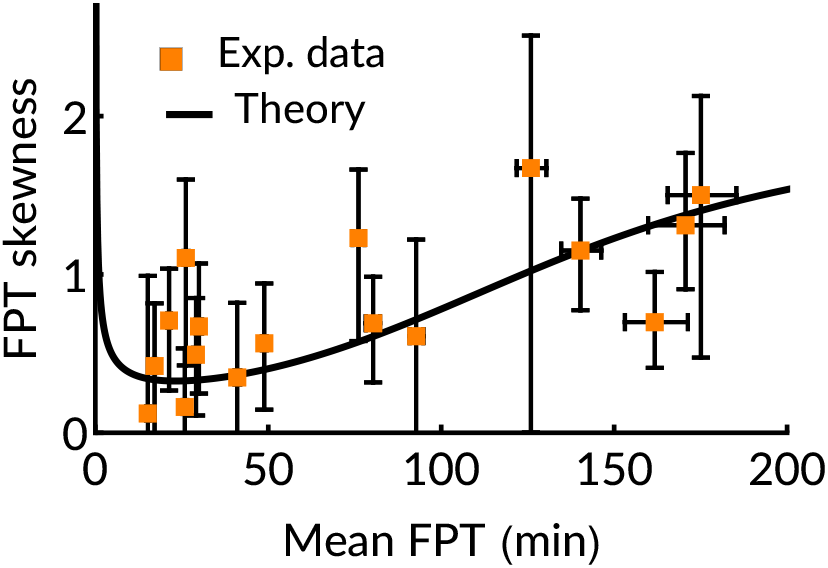
Skewness against ⟨*t*⟩ for 20 mutants (symbols) and exact theory (solid line) with *k* = 4.5 *min*^−1^, *x/b* = 0.01 and *γ* = ln(2)/40 min^−1^. Error bars are determined in a similar way as in Fig.1.

Our results lead to an interesting prediction regarding the dependence of the mean lysis time on the cell doubling time. We showed above that fluctuations are minimal around a mean time *t*_*m*_, for a given cellular size doubling time ln(2)*/γ*. This value *t*_*m*_ for any bacterial cell however may vary, and depends upon experimental conditions which may change the cell doubling time. From the theory, we numerically calculate the *t*_*m*_ at minimum *CV*^2^ and find it to be linearly dependent on the doubling time ln(2)*/γ* (see Fig. 4). We also show analytically that 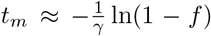 where the fraction *f* is a ratio between an ‘optimal threshold’ *X*_*opt*_ and the steady state concentration *c*_*ss*_ = *kb/γ* (see S8 in SI [35]). Thus, under the plausible assumption that wild-type viruses have optimal noise characteristics, the mean lysis time of lambda phage under different host cell doubling times should be linearly related with cell doubling time. Interestingly, for a wide variety of viruses, a linear correlation between infection delay time and host doubling time has been reported recently [38].

**FIG. 4:**
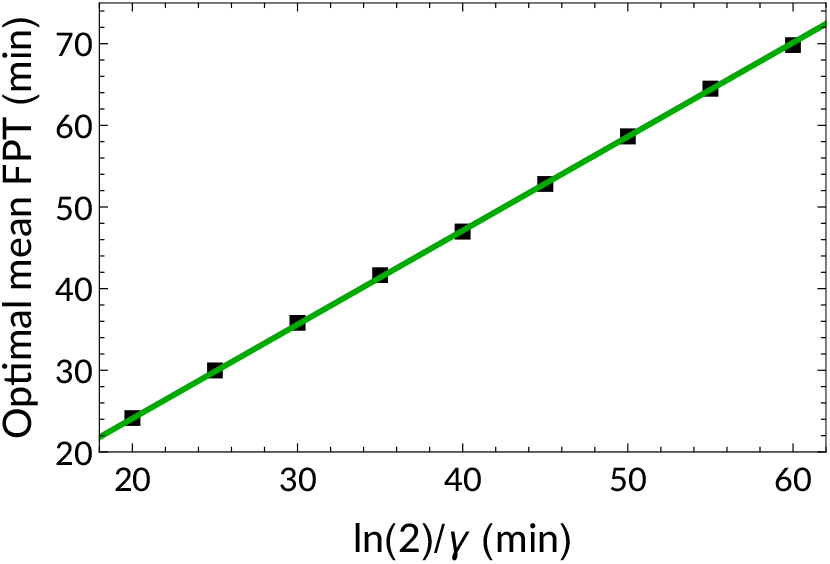
Optimal mean FPT versus cell doubling time. As discussed in the text, slope of the green line is equal to − ln(1 − *f*)/ ln(2) with *f* = 0.55

## Approximate FPTD

Systematic approximations are useful since often they yield simpler expressions of practical use. Previous approximations for moments of the FPTD [20, 24, 39] were based on ad hoc assumptions. We develop a systematic approximate theory which matches the exact results up to second order in fluctuations as follows. Under the assumption that the burst sizes are small, the bursty term in Eq. (1) may be smoothened through the Kramers-Moyal expansion [33] (see S9 in SI [35]). Retaining terms up to second order, we obtain the following backward Fokker-Planck equation, valid for *x* < *kb/γ*:

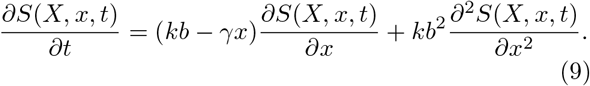

The forward Fokker-Planck equation (S9 in SI [35]) which is the counterpart of the above backward Eq. (9), is the corresponding Kramer-Moyal approximation to the exact theory developed in [7]. That forward equation, under Ito convention, is related to the following Langevin equation [33] (see S9 in SI [35]):

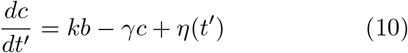

where *t*′ denotes the forward evolving time, and is to be distinguished from the “backward” time *t*. Note *η*(*t*′ is a Gaussian noise with 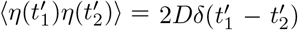, and the diffusion constant needs to be identified as *D* = *kb*^2^. The simple Langevin equation with production, decay, and noise terms have appeared in earlier theories, e.g. of FPT for m-RNA kinetics [40] — yet its exact FPTD was not known. We note that the Eq (9) corresponding to the Langevin Eq. (10), can be exactly solved to obtain the FPTD in Laplace space (see S10 in SI [35]):

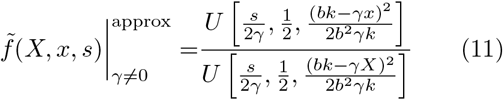

Here, *U* denotes Tricomi’s confluent hypergeometric function related to _1_*F*_1_ (see S10 in SI [35]). From Eq. (11) the theoretical *CV*^2^ and Skewness may be obtained just as we did for the exact theory (Eq. 4) — see S11 of SI [35].

The curves of *CV*^2^ and skewness from the approximate FPTD Eq. (11) are plotted in Fig. 5 along with those from the exact theory (using Eqs. (7), (8)), and the ones from the Langevin simulations obtained using Eq. (10). The exact theory matches the approximate theory and simulations for *CV*^2^ perfectly, but deviates a bit from those in the case of skewness, which is expected since in the Kramers-Moyal approximation we ignored third order moments which contribute to the skewness.

**FIG. 5:**
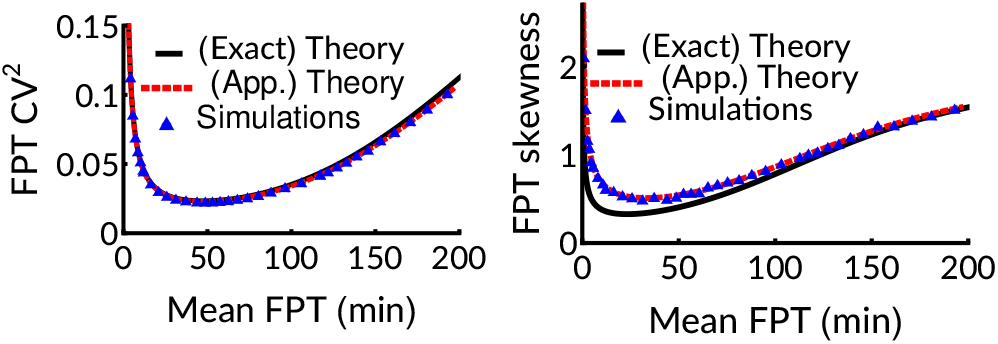
The (a) *CV*^2^ and (b) Skewness, obtained from the exact continuous theory, the approximate theory and the Langevin simulations are compared. The parameters are same as in Figs. (1) & (3).

## Discussion

We derived analytically exact results for the FPTD for protein concentrations by solving the backward Master equation. We showed that the FPTD and its moments match those observed for lysis times in lambda phage, reinforcing the hypothesis that the lysis time is governed by a protein accumulation-dependent FPTD. We developed a systematic approximation for the FPTD and derived analytical results for the approximate FPTD. The results are general and can be applied to fit lysis time distributions for all lytic phages including lambda phage, as well as other protein threshold-crossing processes. The distributions themselves are non-Gaussian with exponential tails and may have a high skewness, indicating asymmetry that may be significant for some lytic phages.

We showed that the FPT theory predicts that some mutants will have optimal noise characteristics, i.e. minimal CV and skewness, which is borne out by the data. The non-monotonic relation between lysis time and noise suggests that lysis time may be selected for during evolution to minimize noise, as suggested by the observation that the wild type lambda phage has the lowest level of noise in lysis times [21].

We also predicted an interesting linear relationship between the optimal mean lysis time and the host cell doubling time. This result may be quite general for lytic viruses, since viral escape from an infected cell is typically characterized by a delay that could be a sign of a threshold phenomena. It is thus intriguing that for a wide variety of viruses, the initial burst timing is linearly correlated with the cell doubling time [38].

## Acknowledgements

D.D. acknowledges SERB India (Grant No. MTR/2019/000341) for financial support. K.R. thanks IIT Bombay for Institute Ph.D. fellowship.

## Supplemental Material for

### S1: Experimental details

Full details of the experimental methods can be found in Ref. [1]. Here we include a brief description in non-specialist language for completeness. Firstly, plasmids were constructed with a mutated version of the holin gene using site directed mutagenesis, by standard molecular biology techniques. These plasmids were then used to transform E. coli lysogens, i.e. cells already harboring a lambda-phage (MC4100 (*λ* cI857 S::Cam) strain) genome. Following induction of the lytic cycle in these lysogens, the mutated holin gene carried by the plasmid is incorporated (via recombination) into the genomes of some of the phages that emerge after cell lysis. The mutated phages were then used to lysogenize wild-type E. coli cells (MC4100 strain), resulting in a collection of lysogenic strains harboring phage genomes with mutations in the holin gene. The phage genome in these lysogens (prophage) expresses a temperature-sensitive cI857 repressor, which is required to maintain the lysogenic state. The lysogen enters the lytic cycle when the cI857 repressor is inactivated following a heat shock. Lysis time was measured as the time taken for a cell to lyse under a microscope starting from the time of application of a heat shock of 42°C. Measurements were carried out with the help of a microscope-mounted, temperature controlled perfusion chamber. Please refer to Ref. [1] for a complete list of the mutant viruses and details of the mutations introduced.

### S2: The continuous Backward Master equation for Survival probability

Let *S*(*X, x, t*) denote the Survival probability of the gene expression process such that the protein concentration stays below threshold concentration *X* at time *t*, starting with concentration *x* at *t* = 0. Since the terminal state (on first passage) of such a process is known (i.e. *X*), the equation governing *S* is developed backward in time *t* and initial position *x*, and the general procedure to construct it is well-known [2]. In this specific problem, between a “ *past* time *t* + *δt*” to a “*future* time *t*” the concentration could have jumped from *x* to *x*′ with probability *kν*(*x*′ −*x*)*δt* due to bursty protein production, where *ν*(*x*′ −*x*) = (1*/b*)*e*^−(*x*′−*x*)*/b*^ is the probability distribution of the burst size *x*′ − *x*. Similarly *x* could have decayed to *x* − *δx* with probability *xγδt*. There is a remaining likelihood that the concentration does not change and stays *x*. Putting these information together one obtains the stochastic Master equation. Note that while the protein production is a jump process, the decay is a smooth process [2], and we are dealing with a continuous variable *x*. Consequently the continuous integro-differential backward Master equation (in the limit *δt, δx* = 0) for the Survival probability is:

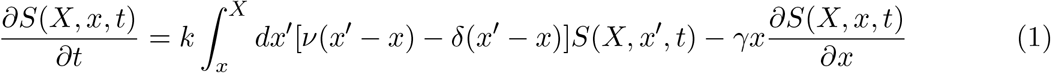

Note that this is the exact counterpart of the forward Master equation of the process developed in [3] for the probability density *p*(*c, t*′) of the protein concentration *c*(*t*′) at time *t*′:

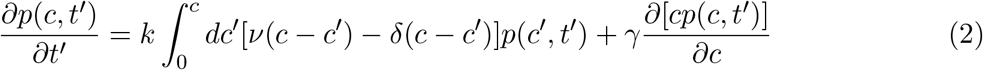

The equation (1) is to be solved for the initial condition *S*(*X, x*, 0) = 1 and the absorbing boundary condition *S*(*X, x* = *X, t*) = 0. We first convert the integro-differential Eq. (1) to a pure differential equation by doing the following mathematical manipulations. Substituting *ν* we have:

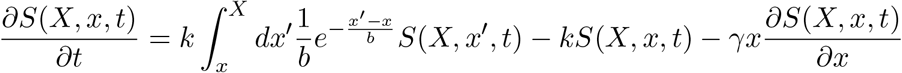

which on integration by parts gives

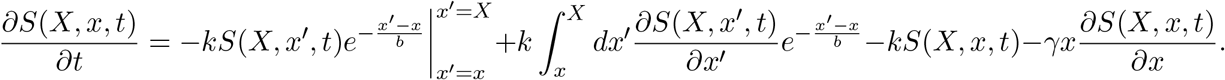

Using the boundary condition, *S*(*X, X, t*) = 0, in the first term,

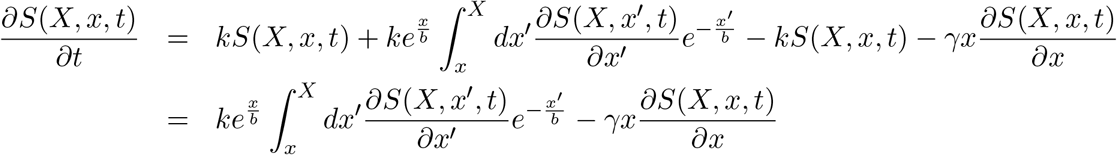

Multiplying by *e*^−*x/b*^, on both sides,

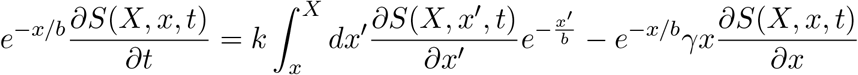

Then we take derivative with respect to *x*, on both sides, and get

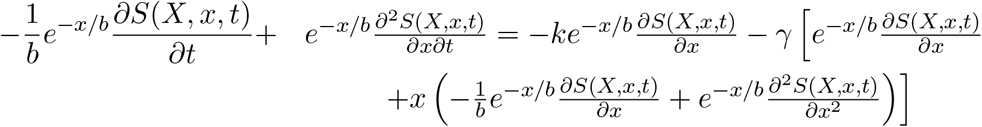

which simplifies to give

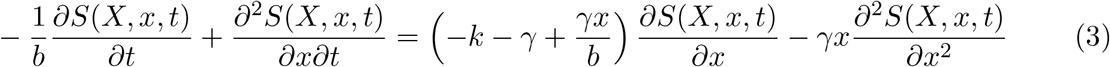

We define the Laplace transform 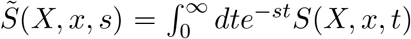, and take Laplace transform of the above equation in time *t* to get,

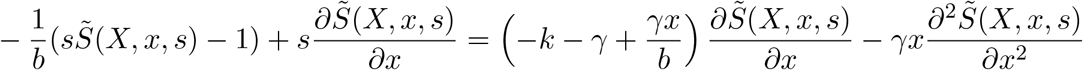

which simplifies to

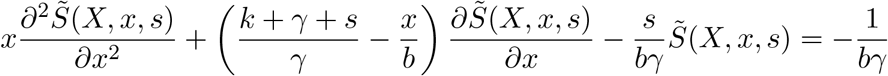

On doing the scale transformation 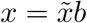, we get an ordinary differential equation for 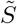:

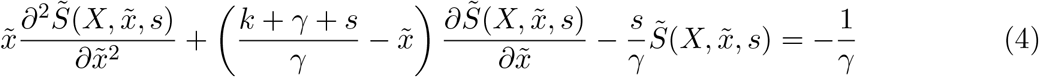

### S3: The exact first passage time distribution (FPTD) in the Laplace domain

The main object of our interest is the first passage time distribution *f* (*X, x, t*) to cross the threshold *X* at time *t*, particularly starting with zero protein concentration (i.e. *x* = 0). This is related to the Survival probability as 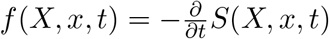. In Laplace space this relationship is 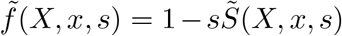. Thus we need to solve for 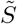 from Eq. (4) to proceed further. The homogeneous part of Eq. (4) is a Confluent Hypergeometric equation, and a particular solution of the inhomogeneous equation is 1*/s* [4]. The general solution is thus:

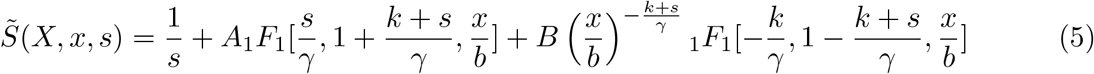

where A and B are constants to be determined and _1_*F*_1_ denotes the Confluent Hypergeometric function of the first kind, 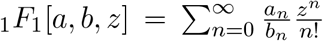, with *a* and *b* being the Pochhammer symbols: *an* = *a*(*a* + 1)(*a* + 2)… (*a* + *n* − 1) and *bn* = *b*(*b* + 1)(*b* + 2)… (*b* + *n* − 1).

The solution is supposed to be valid for *x* = 0. In that limit, the third term in Eq. (5) diverges, because of the factor 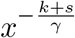. So, we set *B* = 0.

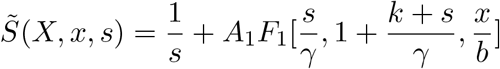

Furthermore using the boundary condition, 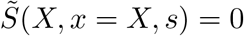, we get the value of *A* to be

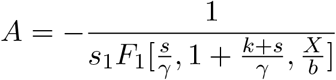

Thus survival probability in Laplace space is

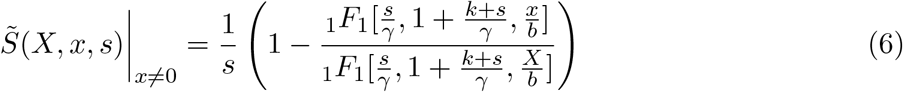

and using the relation 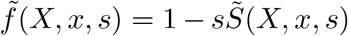, we finally get the central result of this work, namely the FPTD in Laplace space:

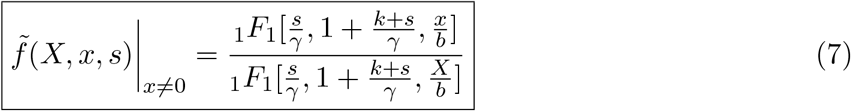

Expressions (6) and (7) are valid for any initial protein concentration *x* > 0. The case of *x* = 0 requires a separate treatment.

#### Solution for the case of *x* = 0

A separate solution for this case is necessary due to the following reason. Consider the case of *x* = 0 and *X* tending to zero. While *S*(*X* = 0^+^, 0, *t*) is something finite, *S*(*X* = 0^+^, *x* = *X, t*) = 0 (boundary condition). Thus the function *S* abruptly changes like a step function and hence the derivative of *S*(*X, x, t*) with respect to *x* will be a Dirac delta function. That would mean that various quantities in the differential Eq. (3) will become singular objects. So, we cannot proceed through this differential equation method for the case *x* = 0. But instead we may now go back to the integral Eq. (1) and study this case separately. In particular we know that, if the boundary *X* goes very close to 0, we expect the survival probability *S*(*X* = 0, 0, *t*) = *e*^−*kt*^ and the FPTD to be *ke*^−*kt*^. That is because a single jump (with rate *k*), will lead to crossing of the infinitesimal threshold. This means that in such a limit the Laplace transforms of the Survival probability and the FPTD are 1/(*k* + *s*) and *k/*(*k* + *s*) respectively. We would see below that the solution of the integro-differential equation for *x* = 0 is perfectly consistent with this expectation. 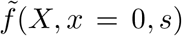 Putting *x* = 0 in Eq. (1) and then taking Laplace transform corresponding to *t*, we get:

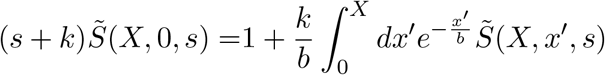

Note that 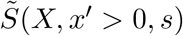 is given by the solution Eq. (6), and hence

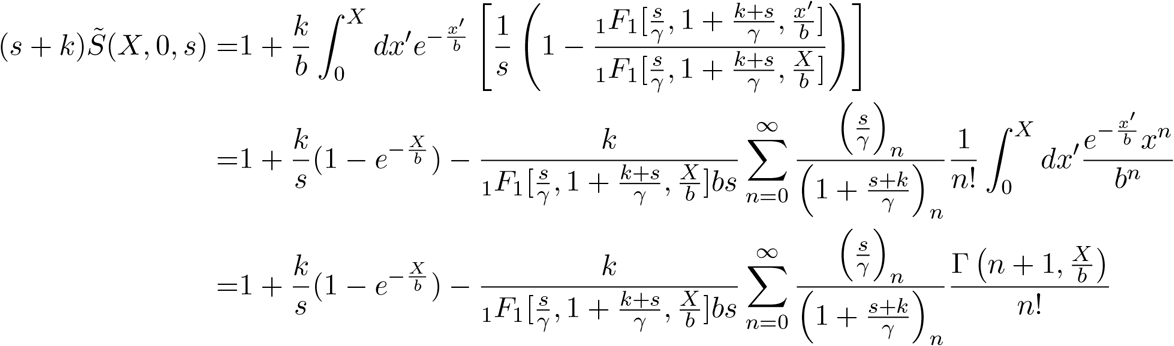

Note the use of the Pochhammer symbols. Thus Survival probability

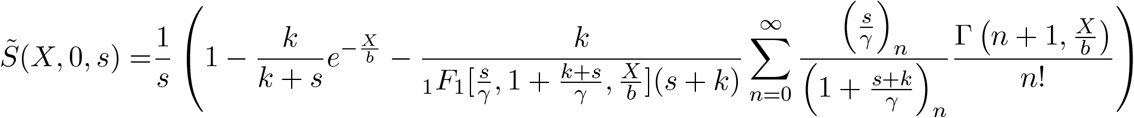

and using the relation 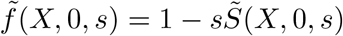, we get,

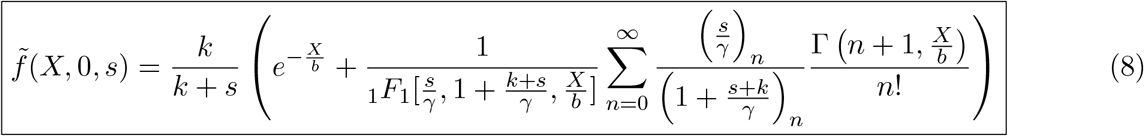

We note that indeed 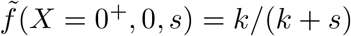 as expected and discussed above.

Although the functional forms of 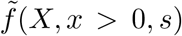 and 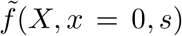 are different (in Eqs. (7) and (8) respectively), as 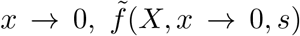 smoothly coincides with 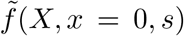. We show this numerically, by plotting the ratio 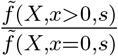, as a function of *x* (see Fig. (1) below). Thus for all practical purposes, the function 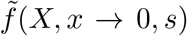 represents well the limit of vanishing protein concentration. Moreover since the function 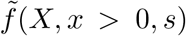 in Eqs. (7) is simpler looking compared to 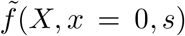, we have subsequently used the expression of 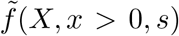 everywhere in the manuscript while discussing vanishingly small initial protein concentration.

### S4: The exact FPTD in the time domain for vanishing degradation and very large cell-doubling times (*γ* = 0)

By writing explicitly the _1_*F*_1_ functions, we have from Eq. (7),

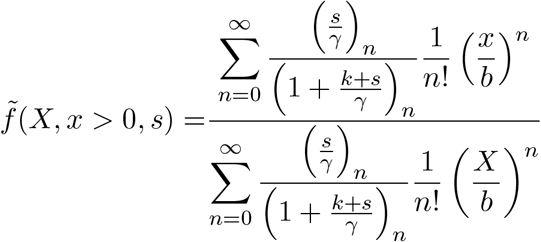

**Figure 1.**
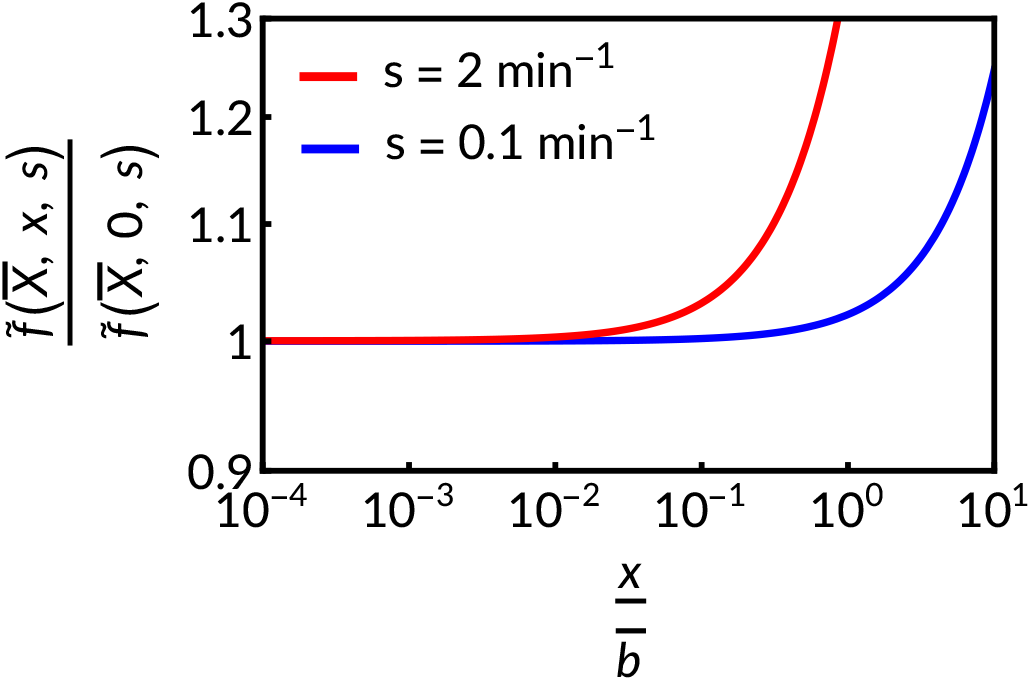
For parameter values *k* = 4.5 min^−1^, *γ* = *ln*(2)/40 min^−1^ and *X/b* = 10, the ratio 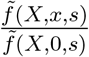 is plotted as a function of *x/b*, for two different values of the Laplace variable: *s* = 0.1 min^−1^ and *s* = 2 min^−1^. In both the cases the ratio approaches 1 smoothly as *x* = 0.

which for *γ* = 0,

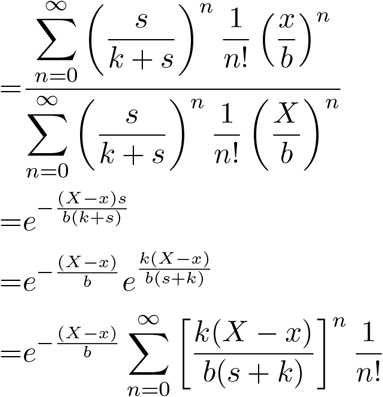

Taking an inverse Laplace transform of the above expression we get,

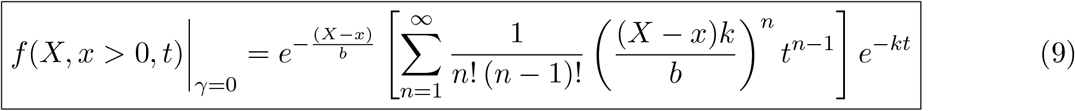

### S5: Moments and cumulants

For convenience of subsequent algebra, we rewrite Eq. (7) as,

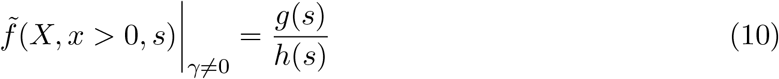

where, 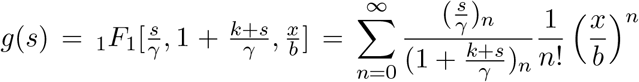 and 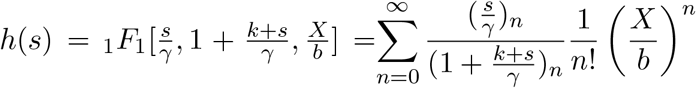.

One of the important use of obtaining FPTD in the Laplace space is that the moments of any finite order may be calculated as derivatives of the Laplace transform at *s* = 0. Namely, the *m*th moment 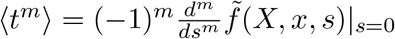. In this section, we calculate the first 3 moments.

#### The mean first passage time (*t* (first moment)

We see from Eq. (10) that

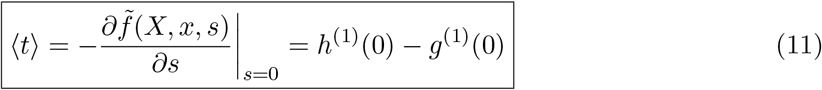

since *g*(0) = *h*(0) = 1, where 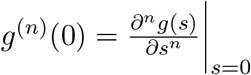 and 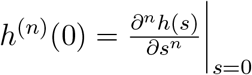. We have

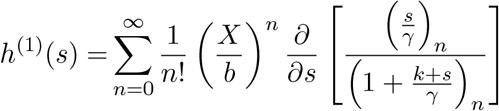

and since for any function 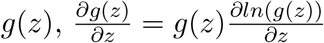, we have

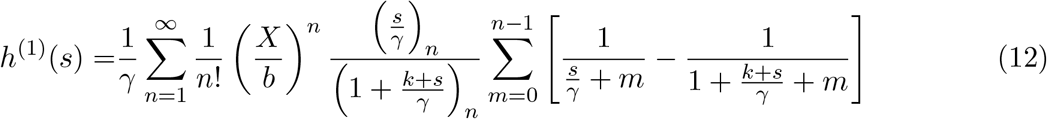

In the limit of *s* = 0, (*s/γ*)_*n*_ = 0. Collecting the surviving terms we have

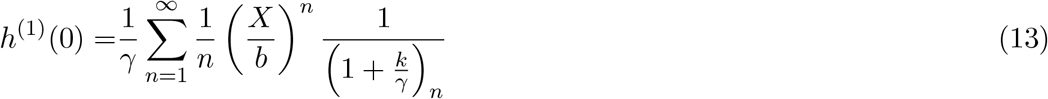

A similar calculation gives

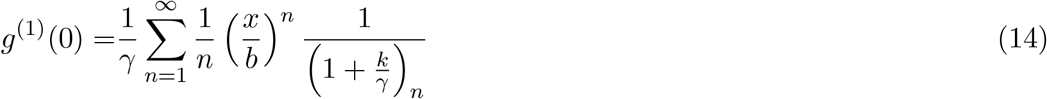

and thus finally

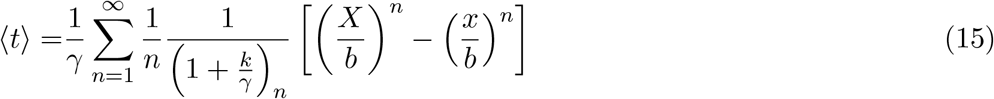

#### The second moment ⟨*t*^2^⟩

Proceeding as above, from Eq. (10),

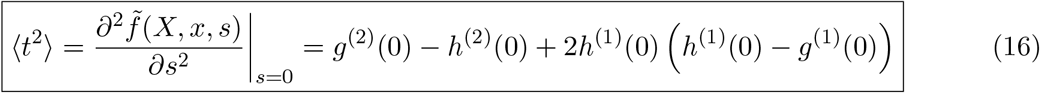

where *h*^(1)^(0) and *g*^(1)^(0) is already evaluated (Eqs. 13 and 14). So, we now need to evaluate *h*^(2)^(0) and *g*^(2)^(0). Differentiating Eq. (12) with respect to *s*:

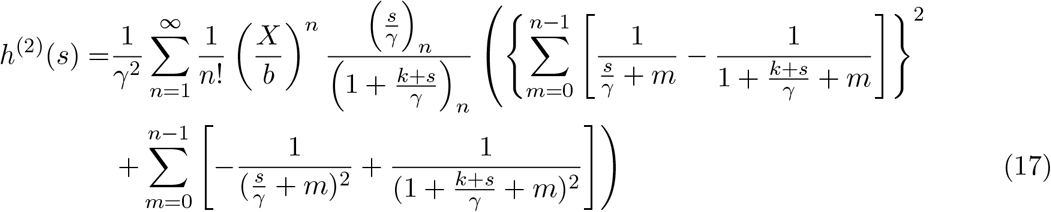

Since for *s* = 0, (*s/γ*)_*n*_ = 0, the non-zero terms yield

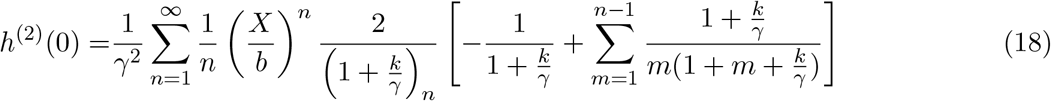

and similarly,

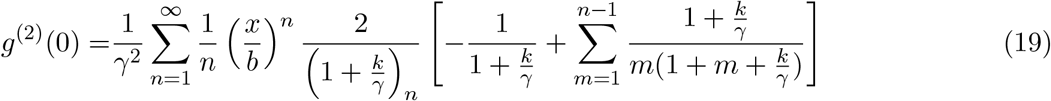

#### The third moment ⟨*t*^3^⟩

From Eq. (10), we get

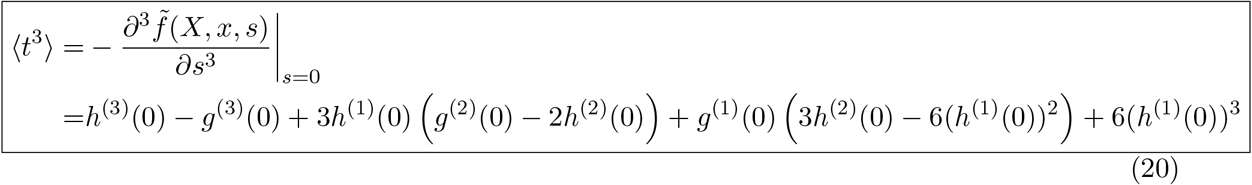

where *h*^(1)^(0), *h*^(2)^(0), *g*^(1)^(0) and *g*^(2)^(0) are given by Eqs. (13, 18, 14 and 19) respectively. So, *h*^(3)^(0) and *g*^(3)^(0) remains to be evaluated. Differentiating Eq. (17) with respect to *s*, we get

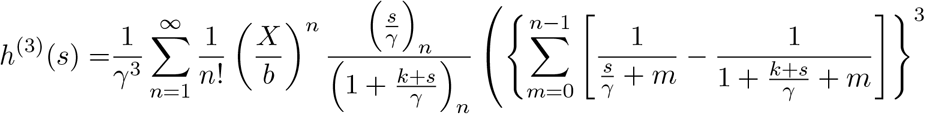

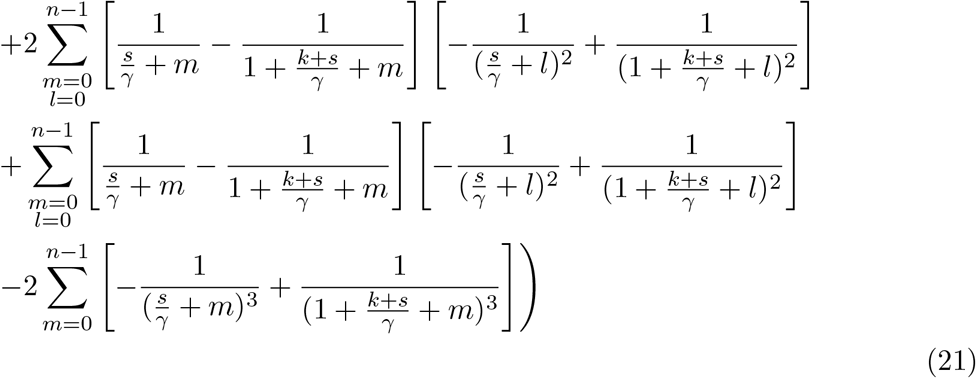

In the limit of *s* = 0, we have

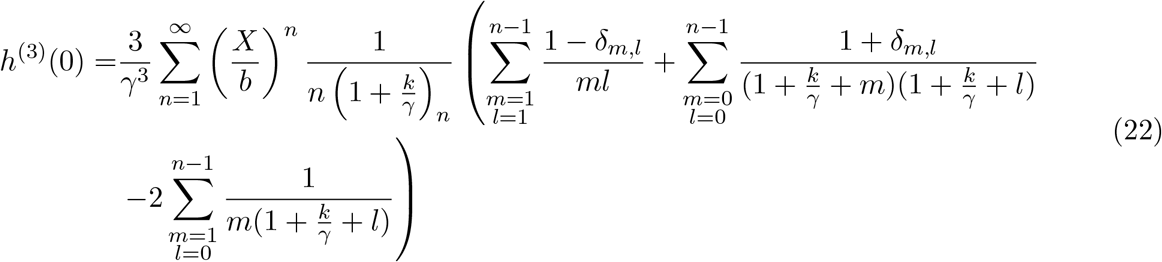

and similarly,

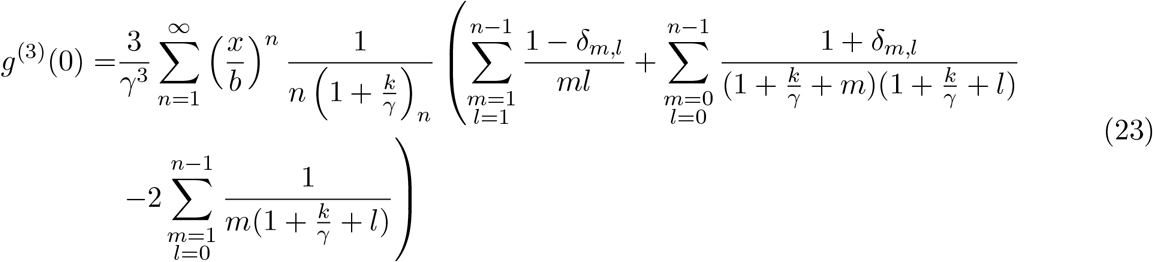

#### The Cumulants

The standard definitions of the *CV*^2^ and skewness are:

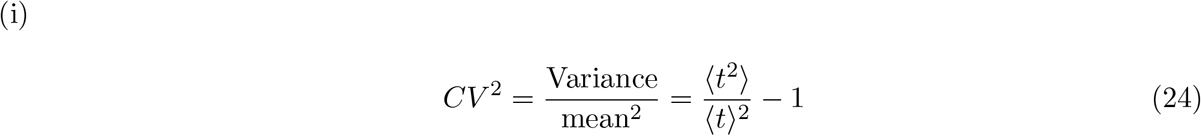

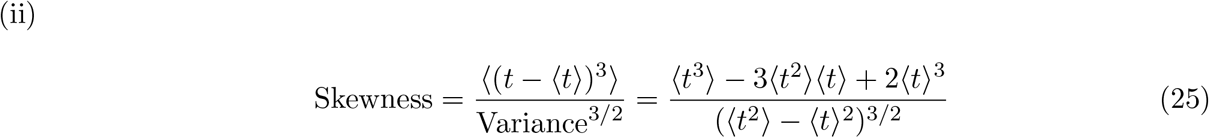

### S6: Fitting of experimental data with our theory to extract the value of *k*

We can calculate experimental values of ⟨*t*⟩ and *CV*^2^ from the 90 - 175 sample lysis times for any of the *λ*-mutants. The theoretical expression of FPTD in Laplace space, and hence all the moments depend on the four parameters-*x/b, k, γ* and *X/b*. We choose *x/b* to be a small value 0.01, as at the beginning of the lytic cycle the protein concentration is supposed to be vanishingly small. E. Coli cell-doubling time is ≈ 40 min (at 37° C) and holin degradation timescale is very large. So, we choose *γ* = *ln*(2)/40 min^−1^. We hope that *k* is similar for all the mutants and they have distinct *X/b*.

We extract *k* by doing a best fit of the theoretical curve of *CV*^2^ versus ⟨*t*⟩ to the experimental data. The procedure is as follows. We vary the value of *k* over a range 1 min^−1^ − 15 min^−1^ in steps of 0.1 min^−1^. For one such value of *k*, using Eq. (15) with upper limit of the sum on *n* being chosen 1000 (a large numerical value as a substitute for ∞), we obtain numerically using Mathematica the value of *X/b*. Using this value of *X/b* and other parameter values as mentioned, using Eq. (16), we obtain a theoretical trial value of *CV*^2^ for that particular mutant. Having 20 such values for 20 mutants, we obtain a sum of squares of the differences of these theoretical trial *CV*^2^ and the corresponding experimental *CV*^2^ of all the mutations. The value of *k* for which this sum of squares is a minimum, gives us the best fit for *k* — we obtained *k* = 4.5 min^−1^.

Once we have this best fitted *k* and the chosen values of *γ* and *x/b*, for any mutant we set the theoretical mean ⟨*t*⟩ equal to the experimental mean FPT, and thus obtain the full time dependent FPTD by numerically inverting the exact FPTD in Laplace space (Eq. (7)).

### S7: Exponential tail and the characteristic time of the FPTD

The FPTD which is the Laplace inverse transform of 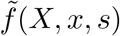 (Eq. (7)) has an asymptotic exponential tail, i.e. for large *t, f* (*X, x, t*) ∼ exp(−*t/τ*_*c*_). The quantity *τ*_*c*_ is called the *characteristic* FPT. It may easily be obtained from a negative pole of 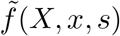as follows. Note that

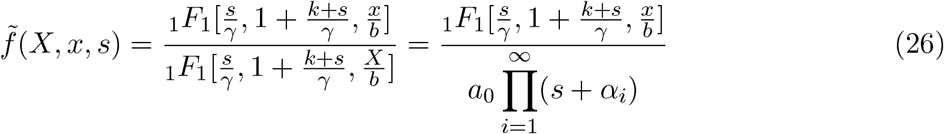

where, −*α*_*i*_’s are the roots of the function 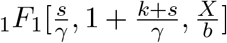 and *a*_0_ is a constant dependent on the parameters. Hence 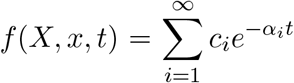 and in the limit of large 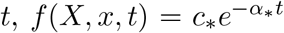

where *α*_*_ is the smallest of {*α*_*i*_}. Thus *τ*_*c*_ = 1*/α*_*_.

For the mutant JD426 with ⟨*t*⟩= 140 min, we first obtain the value of *X/b* numerically solving Eq. (15) for ⟨*t*⟩ = 140 min, *k* = 4.5 min^−1^, *γ* = *ln*(2)/40 min^−1^ and *x/b* = 0.01. We find *X/b* = 241.9. We use this value to find *α*_*_, which is the negative of the root of 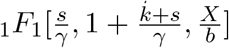 nearest to 0. We get *α*_*_ = 0.0374 min^−1^ and the characteristic first passage time, *τ*_*c*_ = 1*/α*_*_ = 1/0.0374 = 26.72 min. In Fig. (2a), we have plotted the exact theoretical FPTD along with the asymptotic exponential function, 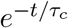, in semi-log scale. The two curves merge for large *t* confirming the exponential tail and the calculated value of *τ*_*c*_.

Next we note that *τ*_*c*_ is a measure of the spread of the FPTD and of fluctuations in FPT. We plot a scaled time *τ*_*c*_/⟨*t*⟩ against mean first passage time ⟨*t*⟩ in Fig. (2b). Like *CV*^2^ and Skewness, we see that it has a non-monotonic shape. For larger mean FPT, the characteristic time gets larger indicating the broadening of the exponential tail.

**Figure 2.**
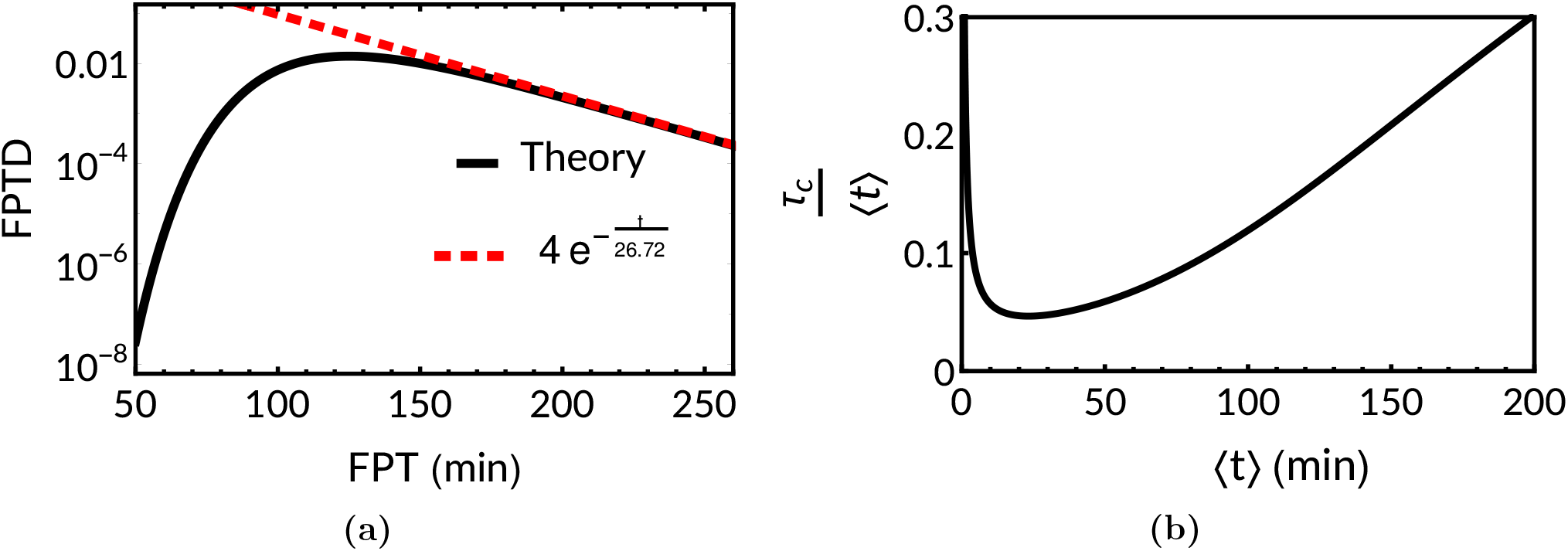
(a) For the mutant JD426, given the parameter values mentioned in the text, the exact theoretical FPTD (solid curve) is plotted (in semi-log scale) along with the asymptotic exponential function (dashed curve) with *τ*_*c*_ = 26.72 min, as a function of the the FPT. (b) Plot of scaled characteristic time *τ*_*c*_/⟨*t*⟩ versus mean FPT ⟨*t*⟩.

### S8: Analytical argument for linear relationship between optimal mean FPT *t*_*m*_ and 1*/γ*

In Eq. (15) above for mean FPT, if we set initial concentration *x* → 0,

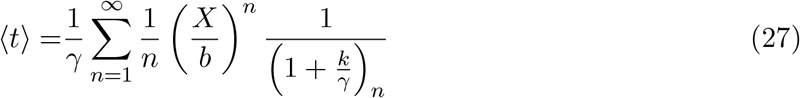

For threshold *X* less than the steady-state mean protein concentration, *c*_*ss*_ = *kb/γ*, in Eq. (27), the term 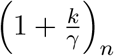 will be clearly greater than (*X/b*)^*n*^. Moreover, 1*/n* term makes the *n*^*th*^ term in the summation decrease fast so that only the first few terms actually contribute. Typically, values of *k* and *γ* are of the order 10^0^ and 10^−2^ respectively. So, *k/γ* will be of the order of 10^2^ ≫ 1. Hence,

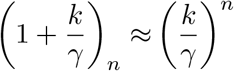

Eq. (27) then becomes

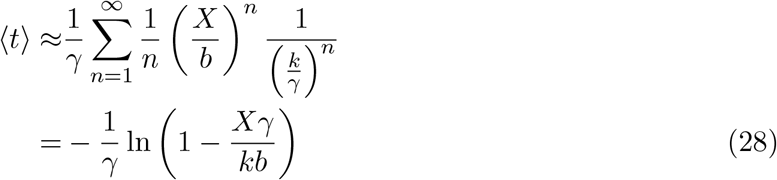

Now if one wants an “optimal ⟨*t*⟩ ” = *t*_*m*_ to minimise fluctuations, one has to minimise the expression of *CV*^2^ (involving the moments in Section S5:) — that calculation does not give simple expressions. Yet one may expect *t*_*m*_ to correspond to some “optimal threshold concentration” *X*_*opt*_, which on general dimensional grounds should be a fraction *f* of the steady state concentration *c*_*ss*_. Thus,

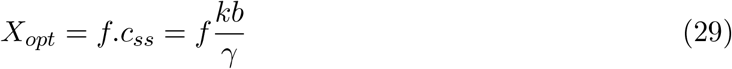

and from Eqs. (28) and (29), putting *X* = *X*_*opt*_ we have optimal mean FPT ⟨*t*⟩ = *t*_*m*_, given by:

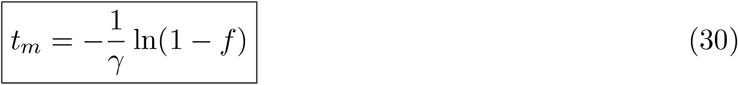

Thus we have shown analytically that a linear relationship is expected between *t*_*m*_ and 1*/γ*. We find the value of *f* ≈ 0.55 numerically.

### S9: Approximate Theory: Fokker-Planck equation and Langevin equation

#### Krammers-Moyal expansion

Krammers–Moyal expansion [2] refers to a Taylor series expansion of the master equation, assuming small jumps *y* = *x*′ − *x*. We first discuss the general procedure to derive a Fokker-Planck equation in this way, and then apply it to the problem of protein production. In general, the backward master equation

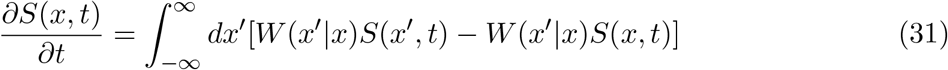

on substituting *y* = *x*^′^ − *x* becomes

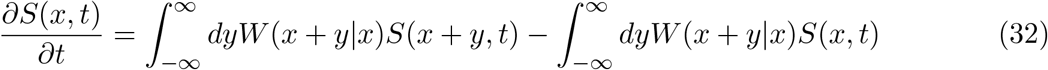

Now, Taylor expanding Eq. (32) around *x*

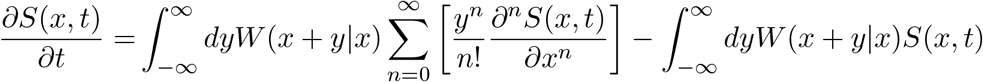

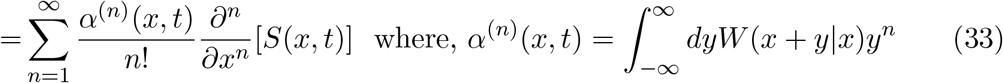

In our problem, the burst size is exponentially distributed, and the burst rate is *k*, such that

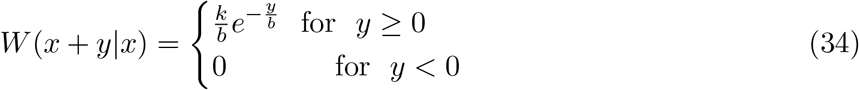

and we may get *α*^(0)^(*x, t*), *α*^(1)^(*x, t*) and *α*^(2)^(*x, t*) from Eq. (33)) as

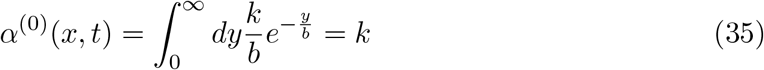

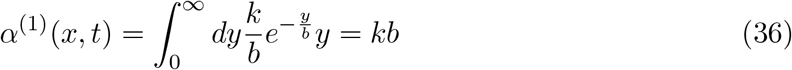

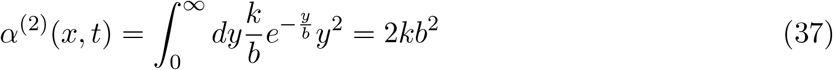

The backward master equation (Eq. (1), including protein degradation, for our problem is

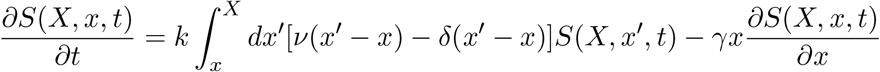

and putting Eqs. (35), (36) and (37) in Eq. (33), and truncating the Kramers-Moyal expansion upto the second-order term we get the approximate backward Fokker-Planck equation as follows:

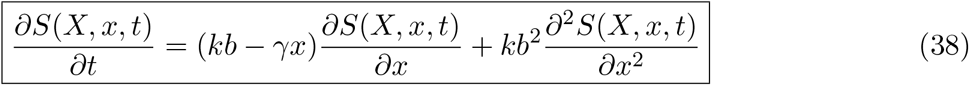

#### The Langevin equation in forward formalism

The forward Fokker-Planck equation corresponding to Eq. (38) follows as a standard result [2,5]:

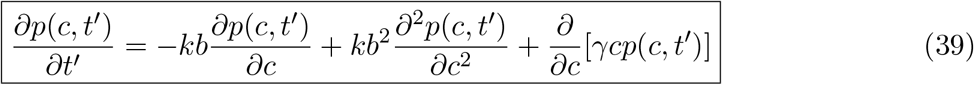

which may be also obtained as the Krammers-Moyal expansion of the forward Master equation Eq. (2) for the protein concentration *p*(*c, t*′) developed in [3]:

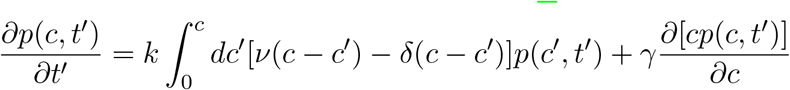

Under Ito’s convention [2,5], the forward Fokker-Planck equation Eq. (39) corresponds to the following Langevin equation:

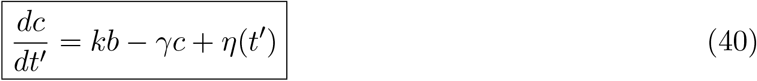

where, *η*(*t*′) is a Gaussian noise with zero mean, ⟨*η*(*t*′)⟩ = 0, and delta correlation, 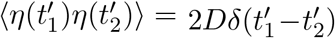, and the diffusion constant needs to be identified as *D* = *kb*^2^. The term *kb* represents production of protein molecules with a fixed burst size *b*. The second term, −*γc*, represents decay of protein concentration.

### S10: Exact FPTD in Laplace space, for the approximate theory

We may exactly solve the Eq. (38) as follows. Defining the Laplace transform 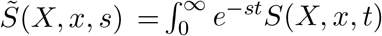, the Laplace transform of Eq. (38) is

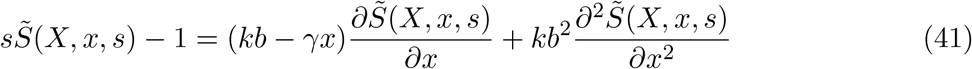

which leads to

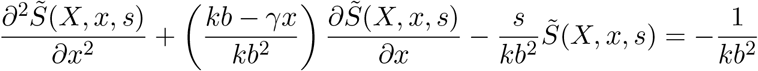

Performing a linear transformation, 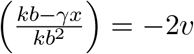, with 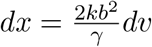 we get

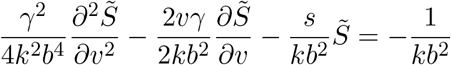

and then a scale transformation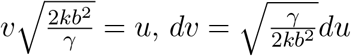 leads to

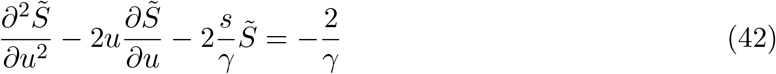

The homogeneous part of Eq. (42) is a Hermite equation [6], and a particular solution of the inhomogeneous equation is 1*/s*. Using the relations between the Hermite function and confluent hypergeometric function of the first kind _1_*F*_1_ and that of Tricomi’s *U* [6], we write the solution of Eq. (42) as:

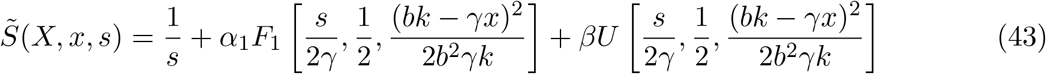

We note that as *k* tends to 0, the term 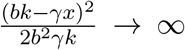, and in such a limit asymptotically, 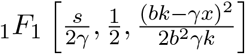 tends to ∞ while 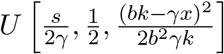 tends to 0 [6]. This would imply that 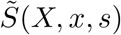 will be unbounded as *k* → 0 for every value of the Laplace variable *s*. Consequently *S*(*X, x, t*) would be unbounded, which cannot be, as it belongs to the range [0, 1]. So, we discard the _1_*F*_1_ solution, i.e., we set *α* = 0.

Using the boundary condition, *S*(*X, X, t*) = 0, we fix *β* and get

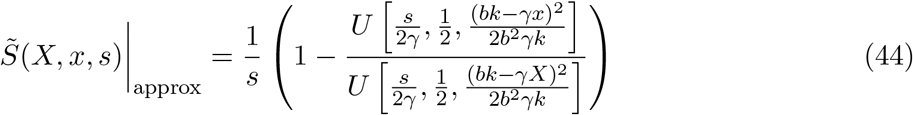

Using the relation 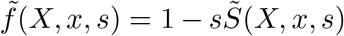, we get the exact FPTD in the Laplace space for this approximate theory:

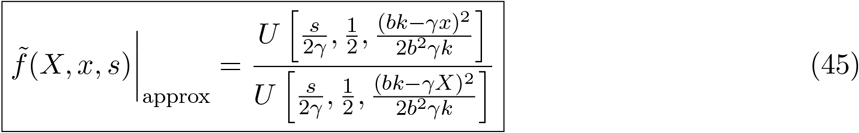

### S11: Exact moments, of the approximate theory

Like Section (S5:) above, we proceed by defining the FPTD (Eq. (45)) of the approximate theory as

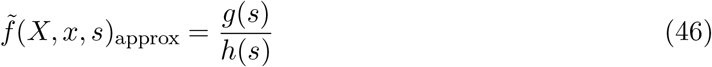

where, 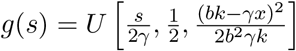 and 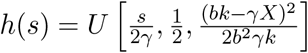.

#### Mean first passage (first moment, ⟨*t* ⟩)

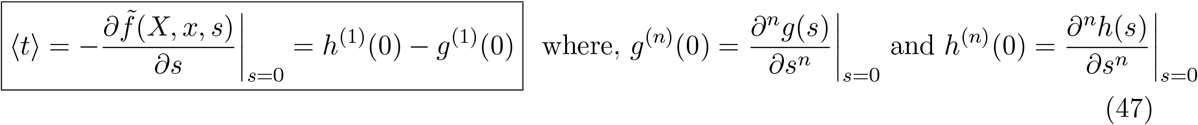

The Tricomi’s confluent hypergeometric function, *U*, can be witten in terms of confluent hypergeometric function (of first kind), _1_*F*_1_, as [6]:

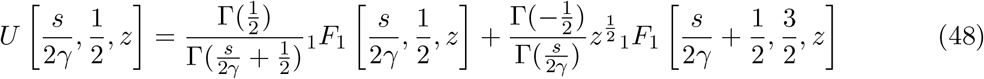

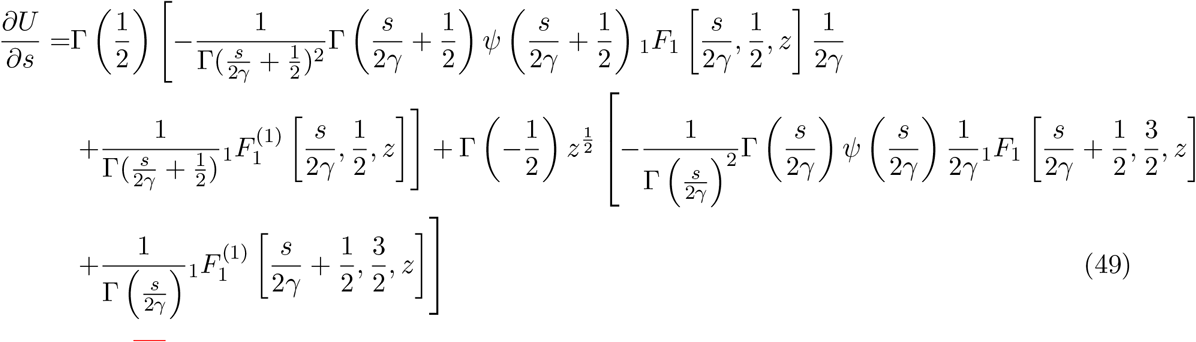

In Eq. (49), the superscipts, (*n*), denotes *n*^th^derivatives and *ψ*(*x*) is the polygamma function defined as 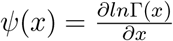 [6]. We note three results (i)-(iii) below.

i. 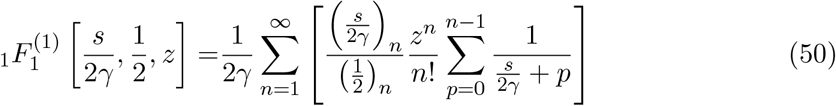

which in the limit of *s* = 0,

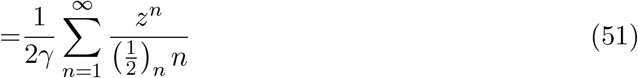
ii. It is know from [6]:

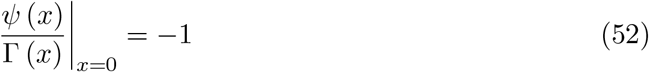
iii. It is known that

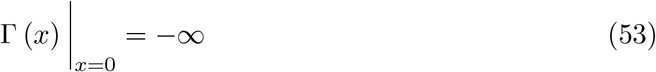

Using Eqs. (51, 52 and 53) in Eq. (49), we get

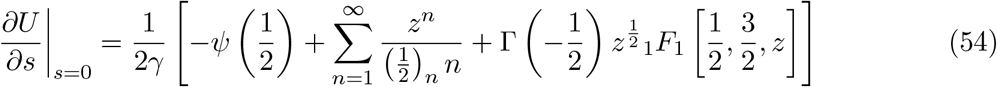

Note *g*^(1)^(0) and *h*^(1)^(0) are given by:

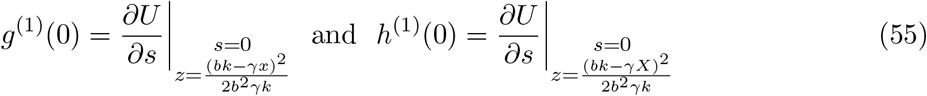

So, using Eq. (54) in Eq. (47), we get the mean first passage time as

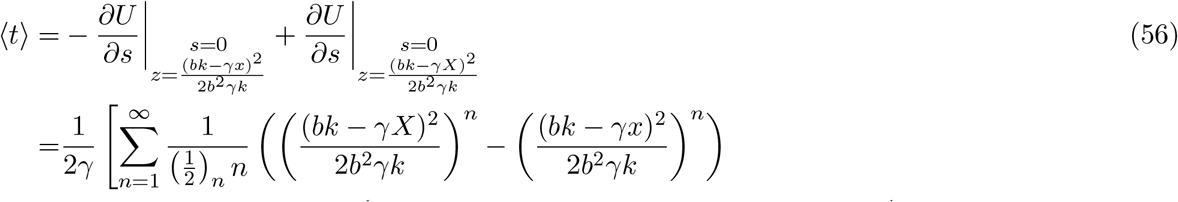

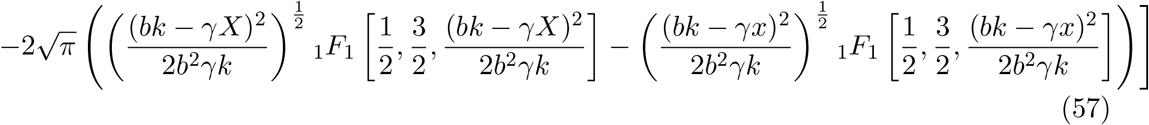

#### The second moment ⟨*t*^2^⟩

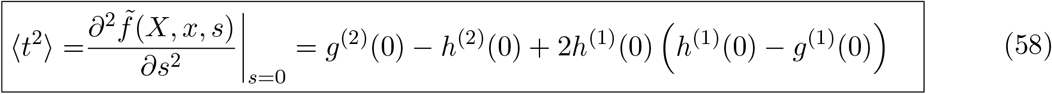

Differentiating Eq. (49) with respect to *s*:

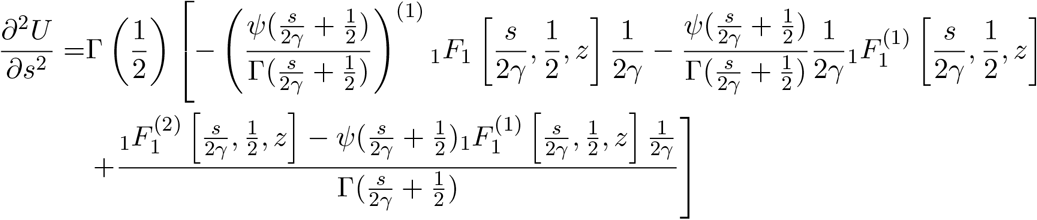

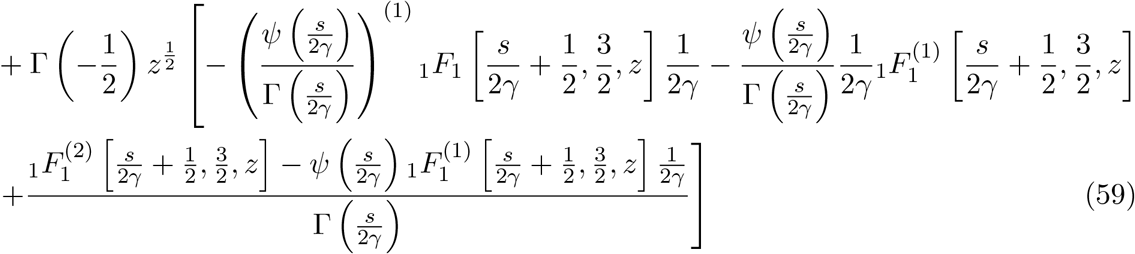

i. From Eq. (50),

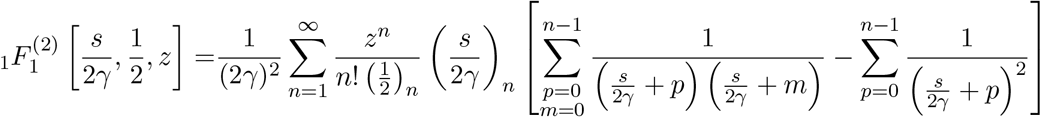

which in the limit of *s* = 0,

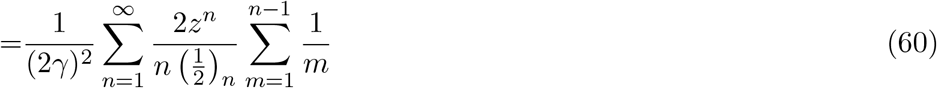
ii. 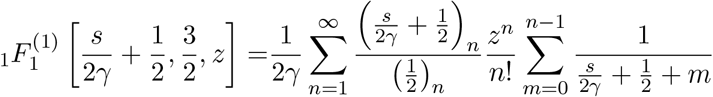

which in the limit of *s* = 0,

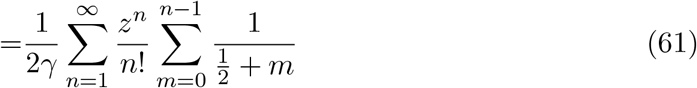
iii. From [6]:

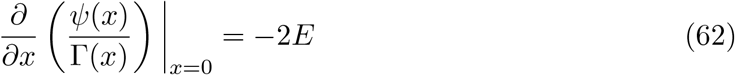

where *E* is Euler’s constant, *E* = 0.57721…

Put Eqs. (51) - (53) and (60) - (62) in Eq. (59),

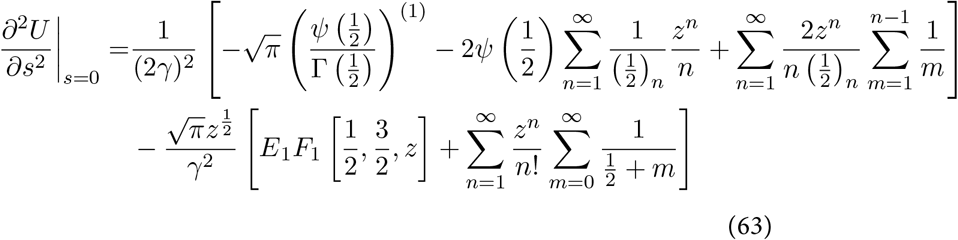

From Eq. (63) we can get the expressions of *g*^(2)^(0) and *h*^(2)^(0):

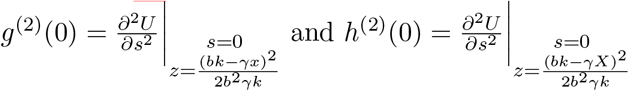

Substituting these expressions along with the previously obtained expressions of *g*^(1)^(0) and *h*^(1)^(0) (Eq. (55)) into Eq. (58), we will finally get the expression for ⟨*t*^2^⟩

#### Third moment, ⟨*t*^3^⟩

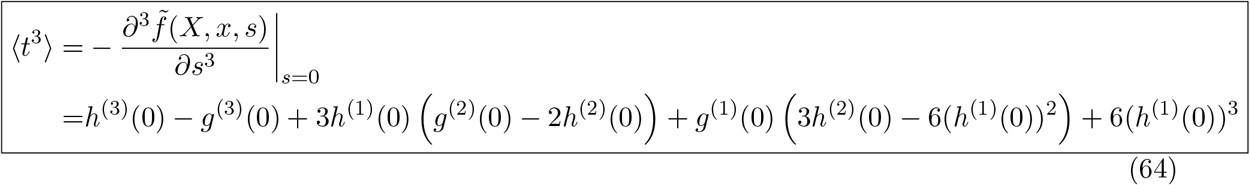

For third moment, we need to evaluate two more terms, *g*^(3)^(0) and *h*^(3)^(0):

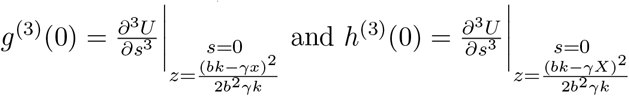

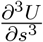 can be ev aluated by taking one more derivative of Eq. (59) with respect to *s*. We skip the calculation as it gets lengthy. The derivatives are easily evaluated numerically by Mathematica which we use for our plots of Skewness in the main manuscript.

